# Long-term ketogenic diet increases expression of SOX2-dependent oligodendrocyte- and myelination-associated genes in the aged mouse brain

**DOI:** 10.1101/2022.02.22.481437

**Authors:** Matthew S. Stratton, Jose Alberto Lopez-Dominguez, Alessandro Canella, Jon J. Ramsey, Gino A. Cortopassi

## Abstract

Aging is associated with multiple neurodegenerative conditions that severely limit quality of life and shorten lifespan. Studies in rodents indicate that in addition to extending lifespan, the ketogenic diet improves cognitive function in aged animals, thus improving healthspan. To broadly investigate what mechanisms might be activated in the brain in response to ketogenic diet, we conducted transcriptome wide analysis on whole brain samples from 13-month-old mice, 26-month-old mice, and 26-month-old mice fed a ketogenic diet. We observed clear activation of inflammation and complement system pathways in the 26-month-old mice relative to the younger animals. Interestingly, ketogenic diet caused a modest but significant increase in the expression of SOX2-dependent oligodendrocyte/myelination markers.

## Introduction

It has long been known that caloric restriction (CR) extends lifespan in model organisms while recent results from human and non-human primate data indicate that, at a minimum, biomarkers of aging and aging-associated morbidity are improved with CR (1–11). Similarly, ketogenic diet (KD) was shown to increase lifespan and healthspan in rodents (12), though the ability of KD to increase lifespan is not without controversy (13). Nonetheless, the ketogenic diet clearly improves cognitive function in aged rodents and decreases inflammation (e.g. 12–15). However, investigation of molecular mechanisms engaged in the brain by KD have largely focused on known mediators of inflammation (e.g. inflammasome) and age-associated changes in mitochondria, with limited use of unbiased approaches. We employed RNA-seq to analyze the whole brain transcriptomes of 13-month-old, 26-month-old, and 26-month-old mice treated with KD. Importantly, the tissue used in these studies was from subject animals of (12), where 14 months of KD treatment resulted in increased longevity, increased physical function, and improved learning/memory scores. Long-term adherence to either the CR or KD regimens in human subjects is challenging and rarely achieved. For instance, in studies of cancer patients enrolled in KD intervention (with an acute reason for adherence), 1-3 month attrition was ~50% (reviewed in 16). The molecular signatures generated here provide a valuable tool for target identification in the quest to prevent cognitive decline in the aging population.

## Results

### The effect of age on gene expression in the mouse brain

Transcriptomics analysis was accomplished on RNA isolated from young (13 months of age) and old (26 months of age) whole brain samples, with both groups being control diet fed. Using strict bioinformatics statistical norms, only 34 genes were significantly different between the two groups. Given the cellular heterogeneity of the brain and the discovery/target identification nature of the experiment, we opted to accept unadjusted p<0.025 as a significantly altered gene. Using minimum expression (baseMean > 0.75) and minimum fold change (log2FC>I.2I) cutoff criteria, this resulted in 247 differentially expressed genes in the young and old brain samples (Figure 1A). A core analysis of these genes was conducted in IPA (Qiagen). Consistent with existing literature, the top effected canonical pathways were related to the complement system and immune system activation (Figure 1B). We also report activated (Figure 1C) and inhibited upstream regulators (Figure 1D), and causal networks (Figure 1E) predicted in the IPA analysis. Notably, lipopolysaccharide (LPS) was the top activated upstream regulator predicted in the analysis and genes leading to this prediction are depicted in a network diagram (Figure 1F).

**Figure 1.**
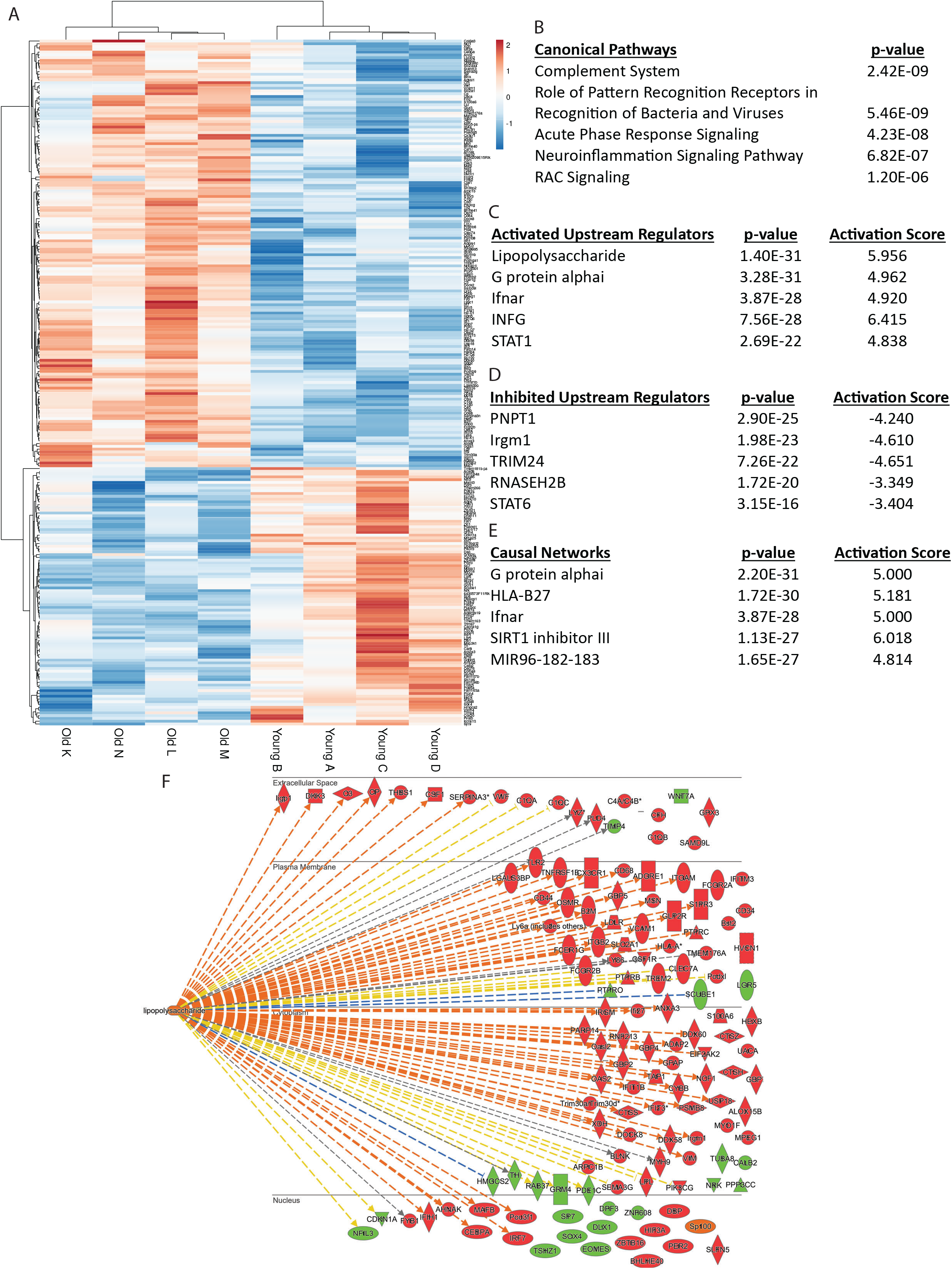
Aging activates pro-inflammatory gene expression patterns in the brain. Differentially expressed genes between 13-month-old and 26-month-old brain samples are shown in heatmap format (A). IPA (Qiagen) was used to determine canonical pathways (B), activated (C) and inhibited (D) upstream regulators, and causal networks (E) predicted to be affected by aging. Lipopolysaccharide (LPS), a pro-inflammatory molecule, was strongly predicted as an activated upstream regulator of the observed gene expression changes in the aged brains. Genes leading to this prediction are shown (F; orange and blue lines indicate measured gene expression is consistent with LPS activation, yellow lines indicate measured gene expression is inconsistent with LPS activation while grey lines indicate LPS is predicted to impact the expression of the target gene without prediction of directionality; red gene symbols indicate increased expression in the aged brain; green gene symbols indicate decreased expression).

### The effect of ketogenic diet on gene expression in the aged mouse brain

Transcriptomics analysis was conducted on brain tissue RNA isolated from young (13-month-old) mice treated with control diet and old (26-month-old) mice treated with KD for 14 months. Again, given rationale above, we opted we to accept unadjusted p<0.025, baseMean > 0.75, and log2FC>I.2I as a significantly altered genes (strict norms only indicated 3 genes significantly altered – Mag, Lgi3, and Serpini1). This analysis resulted in 108 differently expressed genes with KD (Figure 2A). Again, core analysis was accomplished in IPA and we report top effected canonical pathways (Figure 2B) and activated upstream regulators (Figure 2C) predicted in the analysis. Notably, no upstream regulators were predicted to be significantly inhibited with KD, however two causal networks were predicted to be inhibited by KD: BI 836845, an IGF1 neutralizing antibody, and CHD8, a chromatin remodeler (Figure 2D). Both SOX2 and IGF1 were predicted as KD activated upstream regulators and causal network nodes in the analysis. Differentially expressed genes leading to the prediction of SOX2 activation are shown (Figure 2E). Additional IPA based analyses were used to get an indication of KD effects on aging-activated networks and aging effect on KD-activated networks. Specifically, the pathway “build” and “grow” tools allowed analysis of the KD effect on the LPS-dependent genes activated in aging (Supplemental Figure 1). Though KD directly reversed the expression of only a handful of LPS-dependent and aging-induced genes (GRM4, CALB2, PARP14, GBP4, GBP5, Podxl), LPS switched from predicted activation to predicted inhibition with KD and the activity of many of the aging upregulated genes was predicted to be inhibited. Similarly, when the KD-activated and SOX2-dependent genes were compared using the old/young analysis, while the expression of only two genes was reversed (CALB2, GBP4), SOX2 and many of the KD induced genes were predicted to have inhibited activity with aging (Supplemental Figure 2).

**Figure 2.**
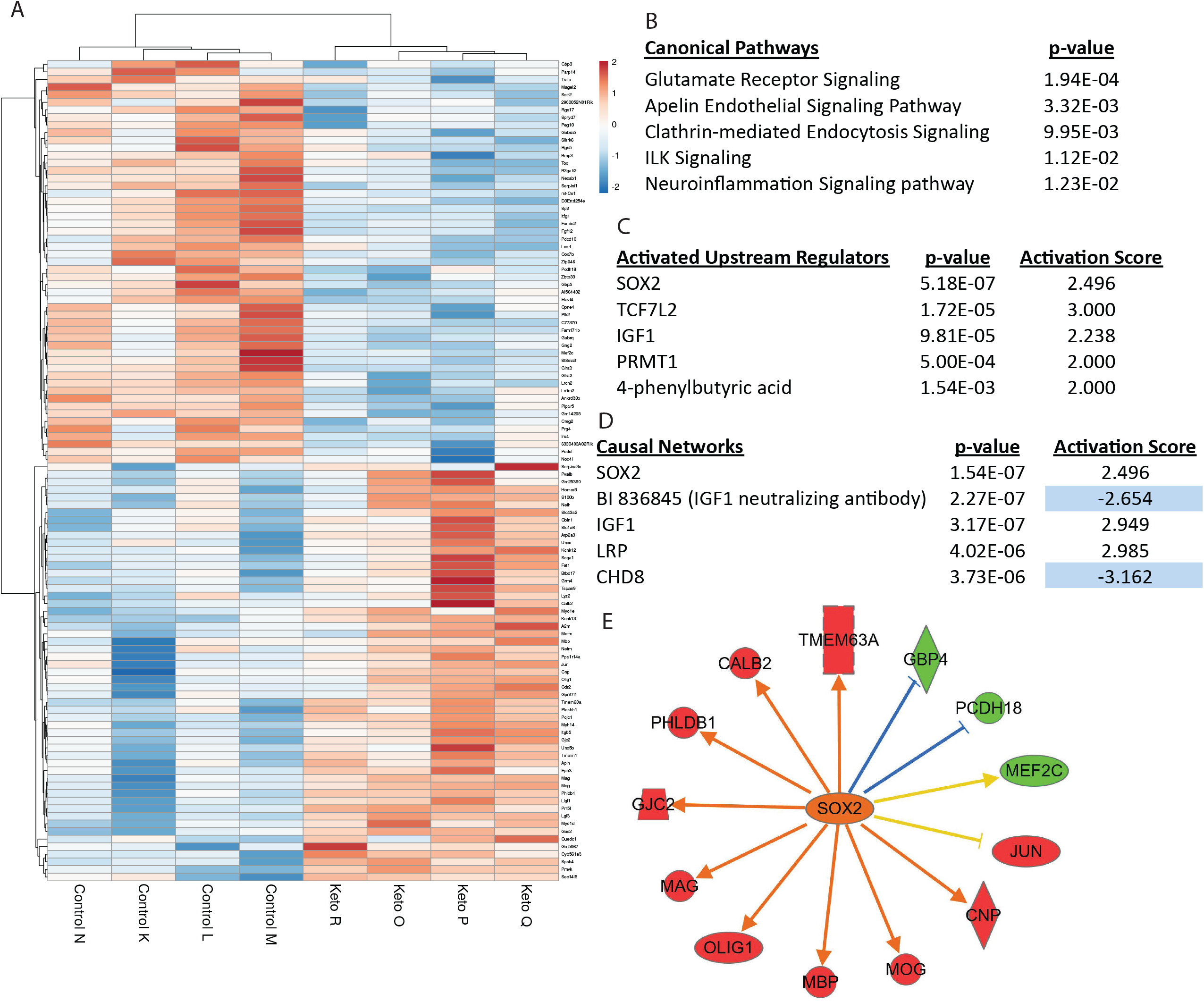
Long-term ketogenic diet is associated with SOX2 activation and increased expression of oligodendrocyte/myelination associated genes in the brain. Differentially expressed genes between 26-month-old animals on a control diet and 26-month-old animals on ketogenic diet are shown in heatmap format (A). IPA (Qiagen) was used to determine canonical pathways (B), activated upstream regulators (C, no candidate upstream regulators were predicted to be significantly inhibited), and causal networks (D) predicted to be affected by ketogenic diet. SOX2, a transcription factor that regulates oligodendrocyte proliferation and differentiation, was the top predicted activated upstream regulator and causal network. Genes leading to this prediction of SOX2 activation are shown, and include a number of myelination/oligodendrocyte specific genes (F; orange and blue lines indicate measured gene expression is consistent with LPS activation, yellow lines indicate measured gene expression is inconsistent with LPS activation while grey lines indicate LPS is predicted to impact the expression of the target gene without prediction of directionality; red gene symbols indicate increased expression in the aged brain; green gene symbols indicate decreased expression).

### Comparing gene expression changes of long-term KD (14 months) with short-term KD in young animals

Remarkably few RNA-seq datasets are available for assessment of KD in the brain and fewer are not on the background of other insult (e.g. epilepsy or brain injury). We selected two RNA-seq datasets where the effect of KD on gene expression in the brain was measured in the absence of other insult. GSE156687 (NCBI GEO) contains RNA-seq analysis from hippocampus of 2-3 month old mice fed control or KD for 3 weeks. Koppel et al., 2021 (17), investigated the effect of 90 days of KD in 4-month old mice. Importantly, this group conducted RNA-seq in neuron vs astrocyte enriched fractions. Differentially expressed genes for Koppel et al were retrieved from supplemental materials. Significantly upregulated and significantly downregulated genes in each of these datasets were used to create gene lists for comparison in GSEA. A log2FC ranked list of our KD dataset was compared to these gene lists using the GSEA “run GSEAPreranked” function. Only significantly upregulated genes in both datasets showed significant enrichment with our ranked list and this enrichment was negative in both instances (Figure 3A and 3B). We next took the genes contained in the leading edges of both analysis (strongest negative association) and analyzed them for pathway enrichment in ConsesnusPathDB (Figure 3C and 3D). There was no overlap between genes in the leading edges or enriched pathways.

**Figure 3.**
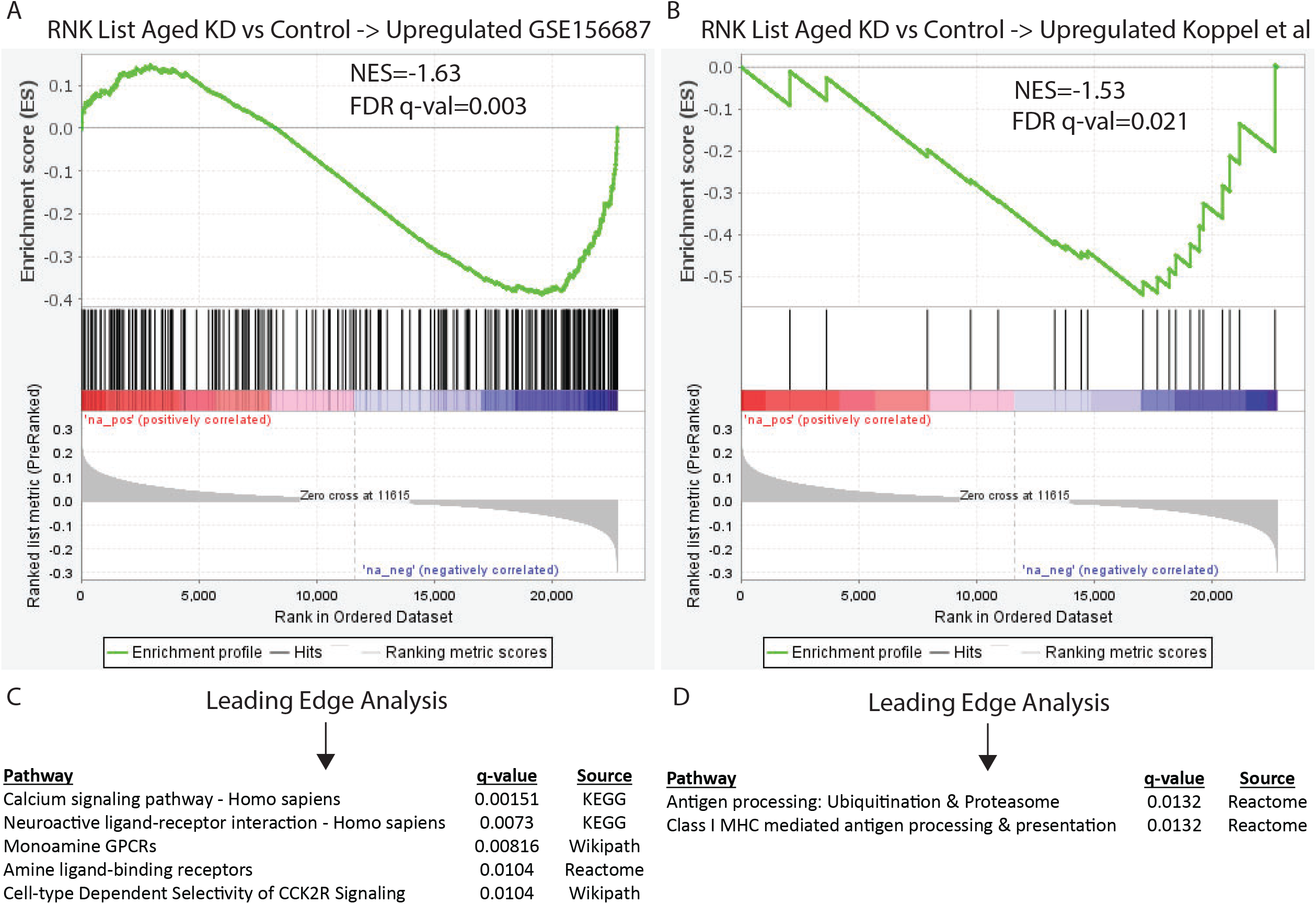
Shorter-term KD in young mice impacts gene expression differently than long-term KD. We identified two KD brain RNA-seq studies where KD was given to young adult mice and used GSEA to compare the effect of shorter-term KD treatment in young adults with the effect of 14 months of KD treatment. Genes found to be upregulated by short-term KD in young animals in both studies were negatively enriched in our long-term KD study (A and B, i.e. a significant portion of the genes activated by short-term KD treatment are inhibited in long-term KD). Direct inspection of the genes contained in the leading edges from both analysis revealed no overlap between the two comparisons. To determine what pathways might be differentially impacted by short term KD in young animals and long-term KD, the genes contained in the leading edge were analyzed with ConsensusPathDB for overrepresented pathways (C and D).

This negative enrichment indicates that some component(s) of the KD effect on the brain is different between short-term treatment in young animals and long-term treatment in older animals. Acyl-lysine posttranslational modifications are know to be upregulated in KD treated and exogenous ketone treated samples. Western blot analysis indicates that this effect remains present in the brain with long-term KD in older animals (Figure 4).

**Figure 4.**
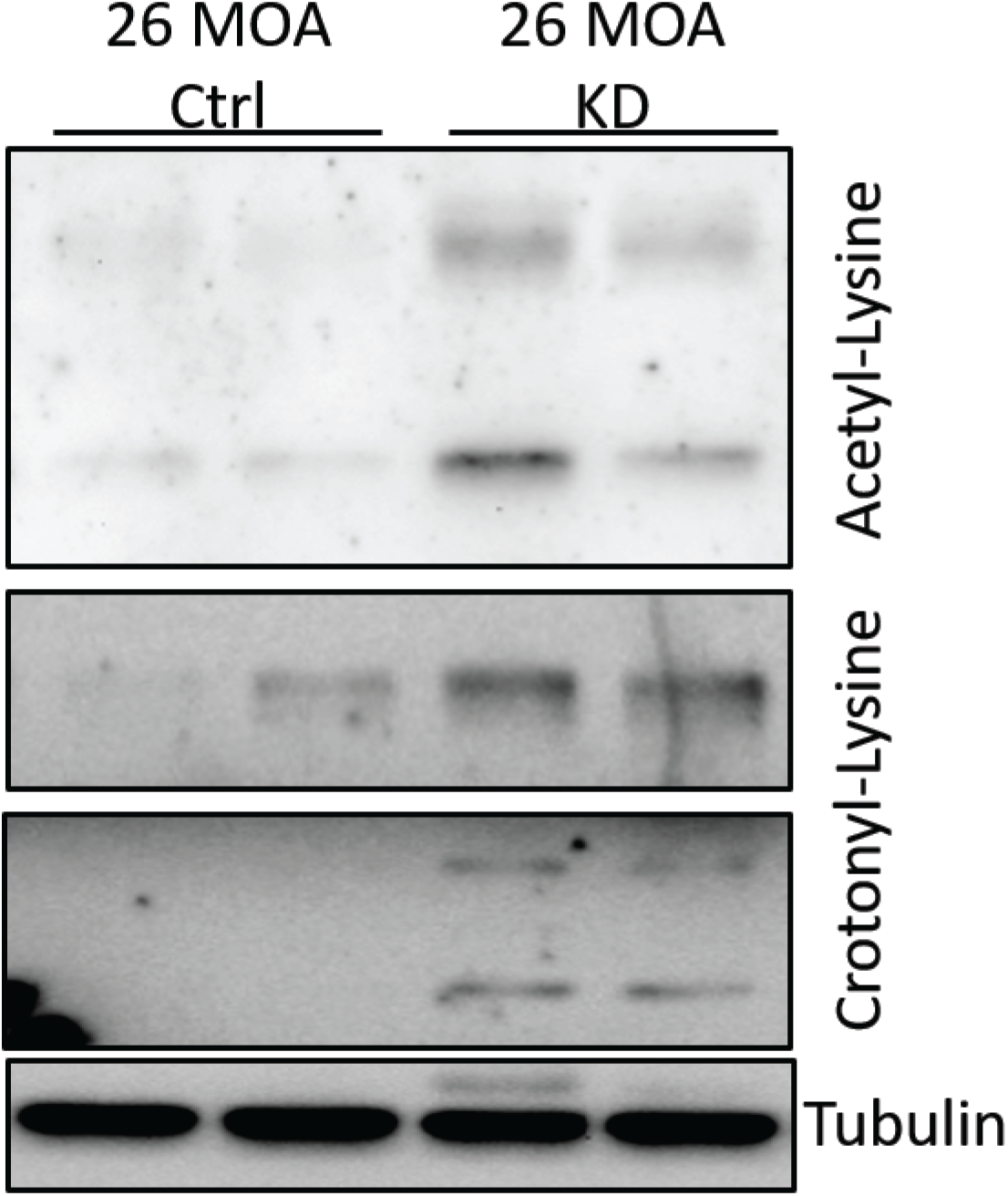
KD increases acyl-lysine protein modifications in the aged brain. The negative association of gene expression changes between shorter-term KD in young mice and KD treatment for ~1/2 the lifespan (Fig. 3) was not expected. We conducted limited biochemical investigation to determine if Western blot probing of acetyl-lysine and crotonyl-lysine modifications in our older/longer KD samples relative to controls indicate that the ability of KD to induced acyl-lysine modifications is independent of age or treatment duration.

## Discussion

Though severe lifestyle modification strategies such as CR and KD show promise for improving brain health in the aging population, long-term adherence to these modifications is difficult. Moreover, the question of how early these interventions need to be employed to achieve meaningful benefit has not been definitively answered. In the pursuit of anti-aging therapeutics, better understanding of mechanisms engaged in brain aging and mechanisms engaged by anti-aging lifestyle modifications is needed. Using an unbiased transcriptomics approach, we demonstrate that 1) aging is broadly associated with activation of complement and inflammatory signaling in the brain, 2) KD activates SOX2-dependent genes that are associated with oligodendrocytes and myelination, and 3) the gene expression effect of short-term KD in young animals is somewhat different from that of long-term KD.

Much of what we identified in these analyses is consistent with the current biology of aging knowledge. For instance, in our old:young analysis dataset, the top predicted inhibited upstream regulator, PNPT1, is a mitochondrial membrane importer of RNA that enables protein translation in the mitochondria. Mutation or genetic targeting of PNPT1 is associated with senescence and accelerated presentation of aging associated pathologies including deafness (18, 19). We also noted an interesting pattern of predicted STAT1 activation and STAT6 inhibition in the upstream regulator analysis. These STATs are know regulators of cell survival in lymphoma (with opposing actions) (20) and have also been shown to have complex roles in immune cell activation (21, 22). Similarly, one of our significantly activated causal networks in the aged brain was for a SIRT1 inhibitor. SIRT1 activation is strongly associated with anti-aging properties (reviewed in 23).

We also report activation of SOX2-dependent genes with long-term KD. A number of these genes are directly involved in axon myelination (Mag, Mog, Olig1, MBP). Given the presence of demyelination in brain aging and the basic function of myelination in neuronal conductance, it is not difficult to imagine how this might be a potentially beneficial mechanism engaged by long-term KD. In fact, KD has shown benefit in multiple genetic demyelination and brain or axon injury models (e.g. 24–26). CHD8, a chromatin remodeler, was identified as the central node for a causal network that is inhibited with long-term KD. Loss of function (mutation or haploinsufficiency) of CHD8 is associated with autism and autism-like phenotypes (27, 28). Finally, our analysis also indicates IGF 1 as an activated upstream regulator of the genes altered by long-term KD. This likely highlights an interesting dichotomy/conundrum in biology of aging research. It is known that long-term inhibition of pro-growth/anabolic signals (e.g. GH and IGF1) slows aging processes, however loss of function in advanced age is at least partially attributed to the age-dependent waning of these same signals (e.g. 29). Would augmenting these signals late in life increase healthspan without shortening lifespan? This remains a burning question in aging research, however data from ketogenic diet in rodents suggests it is possible.

The current study does have limitations. We are unaware of other long-term KD bulk brain RNA-seq datasets. While this highlights the importance of our study, it limits cross-study comparisons. Even short-term KD brain RNA-seq datasets are rare. Of those used for comparison purposes in this study, one was from hippocampus and the other from isolated neuron and astrocyte populations. Conducting RNA-seq on specific cell types in the brain increases specificity of signal but is also subject to artificial gene expression changes of unknown magnitude during the approximately 3hr dissociation and sorting procedure. Along those lines, attempting to compare KD-dependent gene expression changes in young and old mice is complicated by the fact that the aging process changes gene expression and thus, the denominator in the fold change calculation is not constant. Given modest gene expression changes, we loosened the criteria for determining differential expression. We believe this is reasonable given the large cellular heterogeneity of the brain and are encouraged that our findings in whole brain largely reinforced aging biology knowledge and likely represent important pathways that are broadly applicable across multiple brain regions.

## Methods

### Animals

All *in vivo* components of this study were conducted at the University of California Davis using protocols approved by the UC Davis Institutional Animal Care and Use Committee and were in accordance with the NIH guidelines for the Care and Use of Laboratory Animals. All husbandry and dietary intervention was as in Roberts et al, 2017 (12) as samples used in this study were from animals reported on in Roberts et al., 2017. Briefly: C57BL/6 mice were obtained from the NIA aging colony at 11 months of age. After 1 month of acclimation (12 months of age), individually housed animals were randomized to treatment groups of control (18% protein, 65% carbohydrate, and 17% fat as a % of total kcal) or ketogenic diet (10% protein, <1% carbohydrate, 89% fat as a % of total kcal) and fed 11.2 kcal/d. Tissue was harvested after 1 or 14 months on diet, frozen and pulverized (mortar/pestle) with liquid nitrogen.

### RNA isolation

Approximately one-half of the powdered brain was homogenized in 1ml of QIAzol lysis reagent (Qiagen) and RNA was isolated using the Direct-zol RNA Miniprep Plus Kit (Zymo Research), with on-column DNase I treatment.

### RNA-seq

#### Libraries/sequencing

cDNA libraries were prepared with the NEBNext Ultra II Directional RNA Library Prep Kit for Illumina (New England Biolabs). 150bp reads of the cDNA libraries were generated on a NovaSeq SP (Illumina) at the Nationwide Children’s Hospital Genomics Services Laboratory (Columbus, OH).

#### Analysis

RNA-seq analysis was conducted on the Galaxy platform (30, 31). Following trimming using Trimmomatic (32), reads were mapped to the MM10 genome using STAR (33). DESeq2 (34) was used for normalization and differential expression analysis. Heatmaps were generated with Clustvis (35). DESeq2 normalized count and differential expression sheets were loaded to GSEA (Broad Institute, 36, 37) or IPA (Qiagen) for interrogation as indicated in the text. ConcensusPathDB (Max Planck Institute, 38) was used to determine pathway enrichment for leading edge gene sets secondary to GSEA analysis. All RNA-seq data are N=4 male mice per group and no samples were removed from the analysis.

### Western Blot

Approximately one-third of the powdered brain was homogenized in RIPA lysis buffer containing Halt Protease/Phosphatase inhibitors. SDS-PAGE and Western blot procedures were as in (39). Briefly, protein concentration was determined with Pierce BCA assay and 20μg of protein was loaded per well of 10% polyacrylamide gels. Wet transferred nitrocellulose membranes were incubated in anti-crotonyl-lysine (PTM Biolabs), anti-acetyl-lysine (PTM Biolabs), and anti-tubulin antibodies.

## Acknowledgements

We thank Madeleine Lemieux and Patrick Collins for valuable discussion. MS was supported by K01AG056848, AHA857280, and OSU DHLRI startup funding.

## Data Availability

Raw and analyzed sequencing data will be loaded to the NCBI GEO database.

## Figure Legends

**Supplemental Figure 1.**
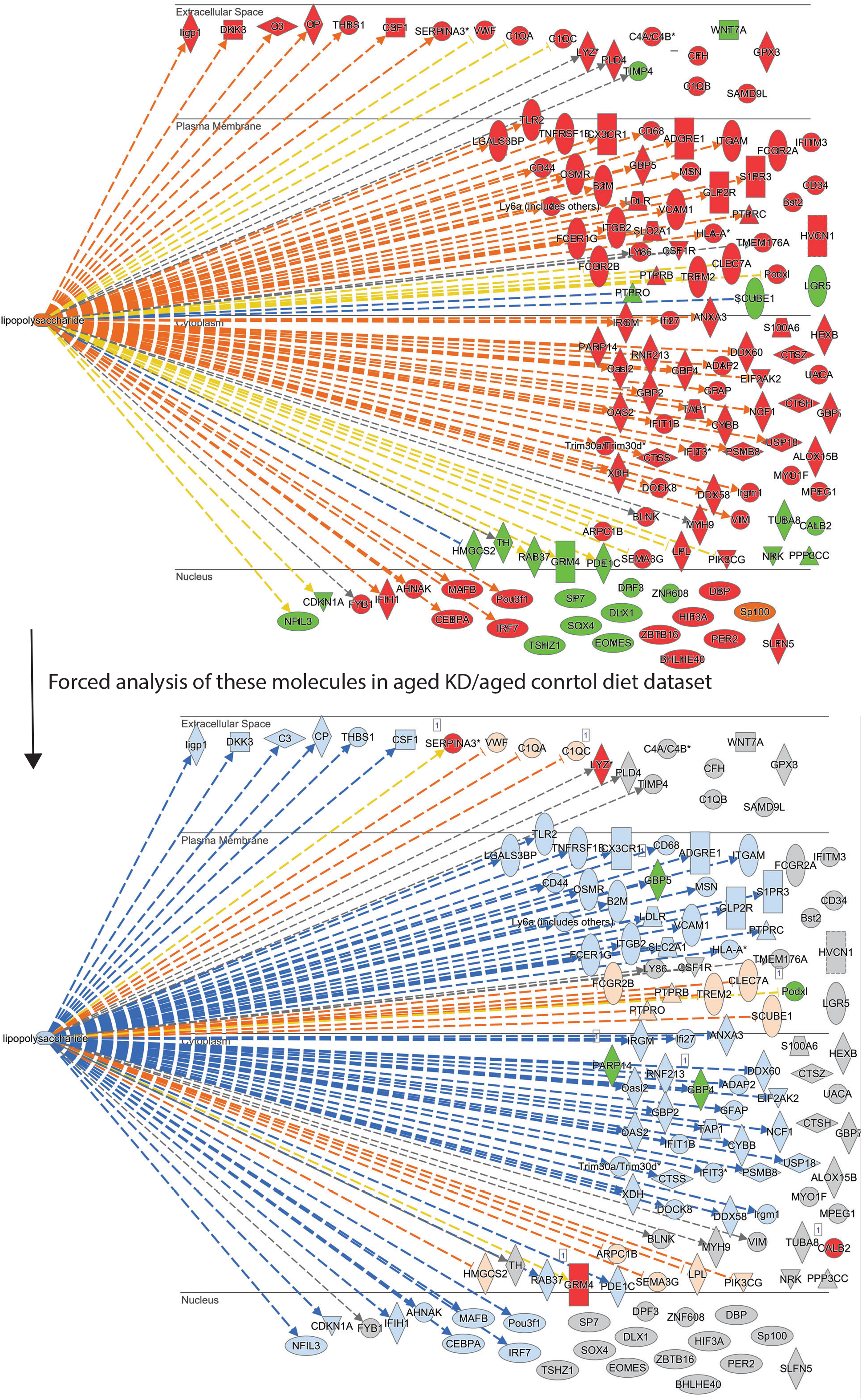
Comparison of LPS activation network from old/young analysis using KD/control data. Using the “build” and “grow” options in the IPA pathway tool, the molecules leading to prediction of LPS activation in the 26-month-old brain samples (Figure 1) were analyzed in the 26-month-old KD to 26-month-old control diet analysis. Though the expression of only a few LPS network genes were significantly impacted by KD, LPS and many of the age/LPS-induced genes were predicted to have inhibited activity (blue gene symbol indicated predicted inhibition, orange indicates predicted activation, grey indicates the gene had neither significantly altered expression nor predicted activity).

**Supplemental Figure 2.**
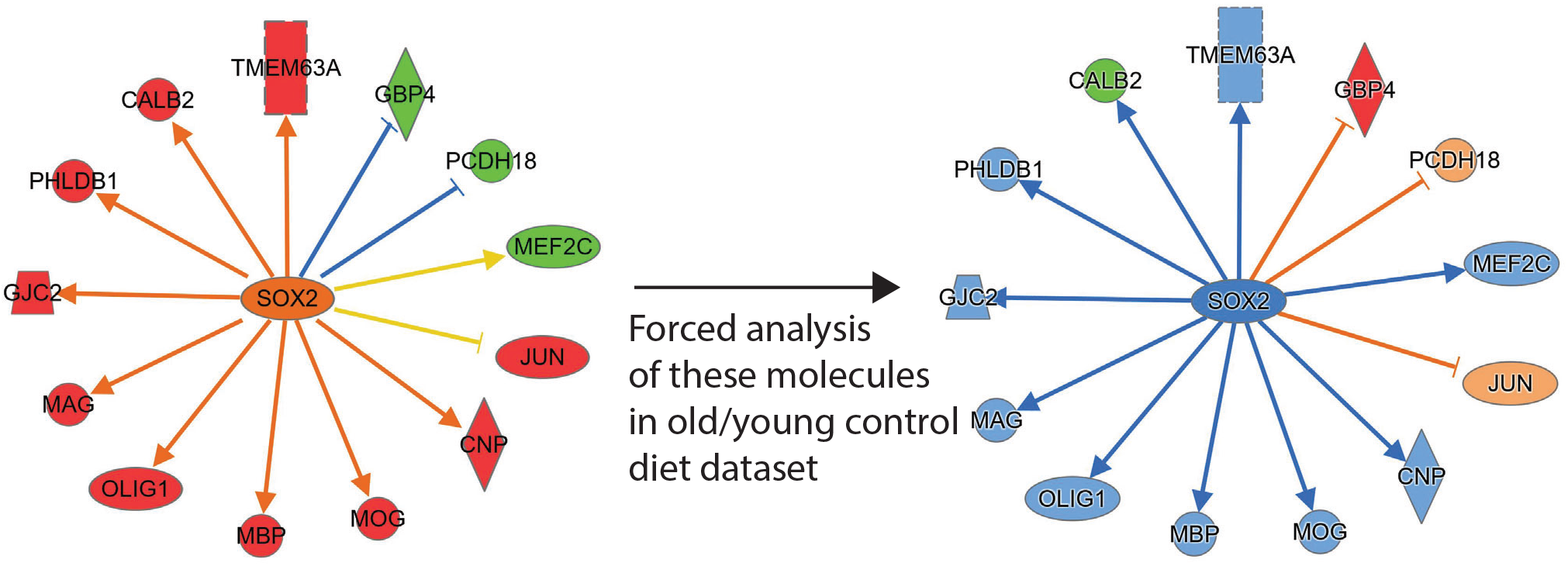
Comparison of SOX2 activation network from KD/control diet analysis using old/young data. Using the “build” and “grow” options in the IPA pathway tool, the molecules leading to prediction of SOX2 activation in the KD samples (Figure 2) were analyzed in the 26-month-old control to 13-month-old control dataset. Similar to supplemental figure 1, though the expression of only two SOX2 network genes was reversed in the comparison, an opposite pattern of predicted activation/inhibition is observed (blue gene symbol indicated predicted inhibition, orange indicates predicted activation, grey indicates the gene had neither significantly altered expression nor predicted activity).

## Notes

### Competing Interest Statement

The authors have declared no competing interest.

